# Phylogenomics of the sigmodontine rodents: Cloud forests and Pliocene extinction explain timing and spread of the radiation of South American mice and rats

**DOI:** 10.1101/2024.12.25.630327

**Authors:** Max R. Bangs, Alexandre R. Percequillo, Víctor Pacheco, Scott J. Steppan

**Affiliations:** Department of Biological Science, Florida State University, Tallahassee, Florida, USA; Departamento de Ciências Biológicas, Escola Superior de Agricultura “Luiz de Queiroz” - Universidade de São Paulo, Piracicaba, São Paulo, Brazil; Museo de Historia Natural, Universidad Nacional Mayor de San Marcos, Lima, Perú

**Keywords:** Great American Biotic Interchange, Oryzomyalia, Target Enrichment, Rodents, Rapid Radiation

## Abstract

Studies of radiations after invasion often overlook the potential role of climatic, biotic, and geologic triggers, instead focusing largely on the conduit for invasion. For example, studies of the rodent subfamily Sigmodontinae, a clade of over 500 species that radiated throughout South America during the Great American Biotic Interchange, have historically focused more on invasion than potential triggers or subsequent environmental change. Here, we put the timing and transitions of this radiation in context of changing climatic, biotic, and geologic factors by reconstructing the biogeography of the radiation. To accomplish this, we generated the largest genomic phylogeny of Sigmodontinae that include over 80% of the genera and 40% of the known species, including all *incertae sedis* taxa and produced a fossil-calibrated chronogram. Results indicate a single invasion of South America at the base of Sigmodontinae (∼ 10.46 million years ago [mya]) with two waves of increased lineage generation and biogeographic transition rates, the first of which occurred after a four-million-year lag following invasion. The timing and location of this initial radiation (6.61 - 5.78 mya) coincided with the spread of montane cloud forest during the Late Miocene Cooling and sigmodontines did not spread throughout the continent until the Mid-Pliocene Faunal Turnover (4.5 – 3.0 mya), a period of high extinction of South American mammals. A comprehensive classification for the subfamily is provided that accounts for the new results.

## Introduction

Causes of increased diversification is a central theme in evolutionary biology; however, most studies have focused on island systems and relatively small clades where the process is more easily studied but are not typical of most biological diversity [1, 2]. Events like the Great American Biotic Interchange (GABI) between North and South America provide an opportunity for radiations to occur across continents, generating more speciose and phenotypically diverse clades that are often understudied due to their diversity of forms (some very rare) and habitats (some quite under sampled) and the difficulties that these issues impose on reconstructing evolutionary and biogeographic histories [see 3].

Muroid rodents are by far the most speciose superfamily in mammals, comprising more than 1,860 species that have diversified across all major continents and island groups [4–6] and as such may yield insights into causes of diversification. Roughly two-thirds of this diversity resides in two clades with unusually high diversification rates, the Old World Murinae and the Neotropical Sigmodontinae (primarily the subclade Oryzomyalia [7]). The strongest case for a radiation within muroids that fit the expectations of an adaptive radiation via ecological opportunity was within the South American members of the Sigmodontinae following colonization by a single species; the Oryzomyalia clade exhibited an early burst followed by a decline in diversification rate consistent with diversity-dependent models, and its members occupied a diversity of habitats and have diverse morphologies [7, 8]. However, other authors have argued for a nonadaptive radiation followed by later transitions into new habitats over time leading to the morphological diversity seen today [9, 10]. Regardless of the mechanism or the precise timing of adaptive divergence, the rapid radiation of this subfamily is exceptional among mammals, generating more than 500 extant species over the last ten million years with species in nearly every ecosystem in South America [5, 7, 11–13].

The rodent subfamily Sigmodontinae represents one of the most recent and diverse mammalian radiations, diversifying throughout South America after crossing from North to South America during the GABI [14] or possibly earlier than this event [15]. However, despite increasing interest in the group [e.g., 7, 10, 13, 16-18], poor resolution of the very short branches at the base of the Oryzomyalia radiation (accounting for approximately 93% of the species; [5]) and limited sampling of the Sigmodontalia sister-clade has led to uncertainty of their place and time of origin (e.g., North, Central, or South American; prior or posterior to the GABI) and relationships among tribes.

Although Sigmodontinae has been the subject of numerous morphological [e.g., 19, 20-23] and molecular systematic studies [e.g., 7, 13, 16, 24, 25-29], these studies have struggled to resolve the relationships near the base of the South American Oryzomyalia radiation. Since 2012, there have been 14 different rooting options of the basal split by molecular phylogenies [9, 10, 13, 17, 27, 28, 30–35], only one of which received strong support: the phylogenomic study of Parada et al. [13] and a subsequent study that combined the Parada et al. [13] data with additional published genes that had much broader taxonomic sampling [18]. Parada et al. [13] produced the first molecular phylogeny with strong support, but they only sampled ∼11% of all known species and ∼40% of all genera, with some tribes and early *incerta sedis* taxa unsampled. Because each of these tribal-level lineages has a distinctive geographic distribution, this limited resolution or sampling at the base of Oryzomyalia has prevented a comprehensive or even accurate reconstruction of the evolutionary and biogeography history of one of the largest mammalian radiations. Vallejos-Garrido et al. [18] added the majority of remaining sigmodontine species, but many of those had very limited genetic data.

While tracing the history of this continental radiation provides many challenges, the advancements in genomic sequencing and availability of continental-scale paleoclimatic reconstructions [e.g., 36, 37, 38] can allow us to tackle this challenge and trace its timing and biogeographic history. In the present contribution, we obtained a comprehensive taxonomic sampling with 74 genera (84% of the 88 known extant genera) and 190 species (∼38% of the >504 known species; [5, 6]) of the subfamily Sigmodontinae, including the critical members of the understudied clade Sigmodontalia and all unplaced *incerta sedis* taxa, allowing us to greatly expand upon the phylogenomic study of Parada et al. [13] and generate the first phylogenomic study sampling nearly all Miocene and Pliocene nodes. Using this phylogeny, we then carried out a time-calibrated biogeographic reconstruction and identified increases in lineage origination and biogeographical transitions in a paleoclimatic context to examine potential triggers for diversification. This study provides one of the largest studies of continental radiation and helps disentangle the complexity of events that shape radiations on this scale.

## Materials and Methods

### Sample selection and library generation

A total of 219 samples were used for this study, representing all five subfamilies of Cricetidae: Cricetinae (3 samples), Arvicolinae (3), Neotominae (9), Tylomyinae (4), and Sigmodontinae (190); as well as Calomyscidae (1) and Muridae (9) as outgroups. Within the New World mice and rats (Neotominae, Tylomyinae, and Sigmodontinae) 16 of 17 tribes were represented; within Sigmodontinae, all remaining tribes and *incertae sedis* taxa (*Abrawayaomys*, *Chinchillula*, *Delomys*) were sampled, excepting the new tribe for the genus *Neomicroxus* [39]. All details on sampling including collection and museum voucher numbers can be found in Supplementary S1 Table.

Of the 219 samples, 180 had Anchored Hybrid Enrichment (AHE) libraries generated using the *Rodent418loci* probe set [40], of which 13 were generated from a previous study across rodents [40]. The remaining 39 samples were from an AHE study focused on the tribe Oryzomyini using the *Vertebrate518loci* probe set [41], that overlapped for 176 loci.

New libraries were generated following the protocol from Bangs and Steppan [40] using either freshly extracted DNA or previously extracted samples from past studies [7, 8, 17]. Additional samples of uncommon species were extracted from either museum skin clips or toe clips using Omega EZNA Tissue DNA Kit following manufacture’s protocol (Omega Bio-tek, Doraville, USA).

All samples were normalized to 1 μg of DNA in 50 μl of Qiagen elution buffer using the broad range Qubit kit to quantify concentrations. Samples were sonicated before library preparation using M220 ultrasonicator (Covaris, UK) for 50 seconds at 50 W peak incident power, 10% duty factor, and 1000 cycles per burst, in order to yield an average size of 500 bps. Some toe clip extractions were already highly degraded with mean size between 500 to 300 bps and were not sonicated but instead moved directly to library preparation. Whole genome library preparation was performed using the NEBNext Ultra II DNA library prep kit (New England BioLabs, UK) and followed the manufacture’s protocol with each individual uniquely indexed using the NEBNext Multiplex Dual Index Oligos for Illumina (New England BioLabs, UK). Libraries were quantified using Qubit broad range kit and pooled at equal concentrations with 24 samples per pool and dehydrated using a SpeedVac vacuum concentrator (ThermoFisher, USA) to produce a concentration of 750 ng of DNA in 3.5 μl of Qiagen elution buffer. Pooled libraries were enriched using the Agilent SureSelect RNA *Rodent418loci* probe enrichment kit following manufacture’s protocol. The final enriched libraries were verified using a TapeStation 2200 (Agilent Technologies, USA) and a qPCR before sequencing on two lanes of Illumina 2500 HiSeq pair-end 150bp sequencing (Illumina, USA) at the GeneWiz (New Jersey, USA).

### Sequence processing and alignment

Reads were processed using the SECAPR pipeline [42], that in brief first removed adaptor sequences, palindromic sequences, and poor-quality reads (Q-score < 20) using Trimmomatic [43], then assembled reads using ABySS [44], before finally pulling out target sequences using a quasi-reference generated using sequences from four Cricetidae samples (*Cricetulus griseus*, *Microtus ochrogaster*, *Peromyscus leucopus*, and *Phyllotis xanthopygus*) from Bangs and Steppan [40]. The reference was a majority-rule consensus of the four samples for all 418 loci and can be accessed at Data Dryad (url XXXX)

Similar to Bangs and Steppan [40], we tested two variables for pulling out target sequences in SECAPR: (1) the percent identity for a match and (2) the minimum coverage of the target on a subset of 24 samples from across the taxonomic breath of the study. Increasing values for these variables decreases the chance for retrieving the correct match, while decreasing these values increases the chance for pulling out multiple matches for a target due to incorrect matches. Values of 40%, 60%, and 80% were tested for the minimum coverage and values of 95%, 90%, 85%, and 80% were tested for the percent identity for a match. Ultimately, we chose 85% percent identity and 60% percent coverage based on maximizing retrieval of target sequences while minimizing loss of sequences due to paralog filtering. Sequences were then aligned using Muscle [45], remapped using a consensus sequence as reference, and finally realigned as suggested by Andermann et al. [42]. All bases with lower than ten read depth were removed using SECAPR in order to exclude potential low-quality areas of the sequences.

All filtered reads have been submitted to NCBI Short Read Archive (BioProject accession number XXX; sample accession numbers XXX - XXX). All alignments, probes sequences, and supporting information is available on Dryad at XXX.

### Phylogenetic analysis

Phylogenies were generated using both concatenated maximum likelihood (IQ-TREE v. 1.6.1; [46]) and multispecies coalescent (Astral-III; [47]) methods. IQ-tree was run using the partition concatenated alignment for GENESITE bootstrap resampling with 1000 ultrafast bootstrap replicates (UFBoot2; [48]) using the CIPRES online portal [49]. This method reduces the chance for overestimation of support for large genomic datasets compared to other maximum likelihood methods [48]. Individual gene trees were estimated in RAxML (v. 8.2.12; [50]) for each AHE locus with 1,000 rapid bootstraps. All nodes with less than 50% bootstrap support were collapsed and resulting trees were used as input into a multispecies coalescent analysis in Astral-III. Collapsing the low support nodes was done to improve accuracy per Zhang et al. [47]. Support in Astral-III was calculated using local posterior probabilities (LPP).

### Fossil calibratation

A time calibrated phylogeny was created using six calibration points used in the most recent Sigmodontinae phylogenic studies [8, 11, 13, 41]. These included one within Muridae (*Mus* – *Rattus* split 11.05 – 12.42 mya *sensu* Kimura et al. [51]; used by Parada et al. [13]), one within Neotominae (*Reithrodontomys* – *Isthmomys* split 1.8-7.49 mya; used by Schenk et al. [8]) and four within Sigmodontinae (Sigmodontini – Ichthyomyini split 4.9 – 14.98 mya used by Schenk et al. [8]) and Parada et al. [13]; *Necromys* – *Thaptomys/Akodon* split 3.5 – 4.1 mya used by Schenk et al. [8]; *Holochilus – Pseudoryzomys* split 0.8-1.2 mya used by Schenk et al. [8]; Schenk and Steppan [11]; and Percequillo et al. [41]; and *Loxodontomys/Auliscomys* 3.5 – 4.1 mya *sensu* Barbière et al. [52] and used by Parada et al. [41]). The only difference in placement of these calibration points from past studies was in the use of the *Kraglievichimys formosus* fossil as the crown age of the clade containing *Loxodontomys/Auliscomys* within Phyllotini. Schenk et al. [8] applied a conservative age estimate of 4.0 – 6.8 mya to the most recent common ancestor (MRCA) of *Auliscomys* and *Andalgalomys,* that is several short branches deeper within Phyllotini, whereas Parada et al. [13] applied this fossil even deeper to the crown age of Phyllotini as a whole, using a narrower confidence interval (CI) of 3.5 – 4.1 mya, with the node-attribution based on a reevaluation of the fossil by Barbière et al. [52]. Barbière et al. [52] concluded that *K. formosus* possessed characters of both *Loxodontomys* and *Auliscomys*, however a well-supported sister grouping of these two genera had never been recovered. Here, however, these two genera were recovered as sister taxa with a high support and thus we apply this calibration to the MRCA of *Loxodontomys* and *Auliscomys*.

Fossil calibration was performed using a full maximum likelihood model in MEGA (v. 11.0.9, Tamura et al. 2021) on a subset of 50 loci. Both recent phylogenomic studies utilizing target capture (ultra-conserved elements [13]; and AHE [41]) used a subset of the loci due to computation limitation and concerns with topological and rate heterogeneity across loci (per Smith et al. [53]). To select the subset of AHE loci we follow guideline of Percequillo et al. [41] for selecting the 50 most clock-like loci using SortaDate [53].

### Biogeographic reconstruction

Species were placed into one or more of ten regions with the ranges of each species extracted from IUCN [54]. A species is considered present in a region if more than 10% of its distribution range is within that region. Regions were delimited based on vegetation, precipitation, and elevation maps and largely follows the Level I ecoregions outlined by Griffith et al. [55]. Using this coding, an ancestral biogeographic reconstruction was performed in RASP using a full hierarchical Bayesian Binary MCMC (BBM) analysis with a F81 + G model with a maximum number of three areas run across ten chains for 50,000 generations, a 1,000 generation burn-in, and sampled every 100 generations. We utilized this method because it has been shown to work well for phylogenies with large number of taxa and allows for (1) differences in transition rates between regions, (2) multi-state reconstructions, (3) samples coded for multiple regions, and (4) uneven distributions across regions (see, [11]).

### New zoological taxonomic names

The electronic edition of this article conforms to the requirements of the amended International Code of Zoological Nomenclature, and hence the new names contained herein are available under that Code from the electronic edition of this article. This published work and the nomenclatural acts it contains have been registered in ZooBank, the online registration system for the ICZN. The ZooBank LSIDs (Life Science Identifiers) can be resolved and the associated information viewed through any standard web browser by appending the LSID to the prefix “http://zoobank.org/”. The LSID for this publication is: urn:lsid:zoobank.org:pub: XXXXXXX.

The electronic edition of this work was published in a journal with an ISSN, and has been archived and is available from the following digital repositories: LOCKSS and FSU Research Repository.

## Results

Samples enriched using the *Rodent418Loci* AHE probe set generated sequences for 415 of the 418 target loci (99.3%) with an average of 405.7 (97.7%) sequences generated per sample and a range of 375 (90.4%) to 415 (100%) sequences per sample. For the additional sequences from 39 samples generated with the *Vert512Loci* AHE probe set [41], 157 loci overlapped and successfully aligned with the 415 loci newly generated for a total sequence overlap of 256,010 base pairs (bps) between the two data sets. The final combined alignment consisted of 538,288 bps of which 164,676 were parsimony-informative and 86,414 were singleton sites, with on average 321,357 bps per sample. All samples had at minimum 150,000 bps of aligned sequence. The alignment for the time-calibrated tree consisted of 50 loci with combined sequence length of 76,859 bps with all samples having at least 32 loci and on average 51,263 bps of sequence. Both alignments can be found on Data Dryad (link to be provided).

The topology of phylogenies generated in IQ-Tree and Astral-III were similar and had similar support values throughout. Most nodes (92%) were recovered with high support (>0.90 UF bootstrap and LPP; Fig 1), with only ten nodes with moderate support (0.89 – 0.75 UF bootstrap and 0.89 – 0.75 LPP) and eight with low or conflicting support (< 0.75 UF bootstrap or < 0.75 LPP). Most of these moderate to low supported nodes were within tribes, at the species or genus level (16 out of 18). The exceptions were the node for the placement of Reithrodontini having moderate support (0.79 UF bootstrap and 0.87 LPP) and the position of tribe Andinomyini having high support but conflicting placement, with either (i) IQ-Tree supporting its placement as the sister taxon to the clade formed by Euneomyini, *Chinchillula*, Abrotrichini, Wiedomyini, *Delomys*, and Phyllotini (Fig 1), or (ii) Astral-III placing Andinomyini within that group sister to a clade formed by Euneomyini and *Chinchillula*. The placement of Andinomyini had no effect on the biogeographic reconstruction given that the three nodes leading up the split as well as the one after the split all reconstructed to the same region and also represents one of the shortest internode distances in the phylogeny with the fossil calibrated phylogeny estimating the duration at ∼63,700 years (Fig 2).

**Fig 1:**
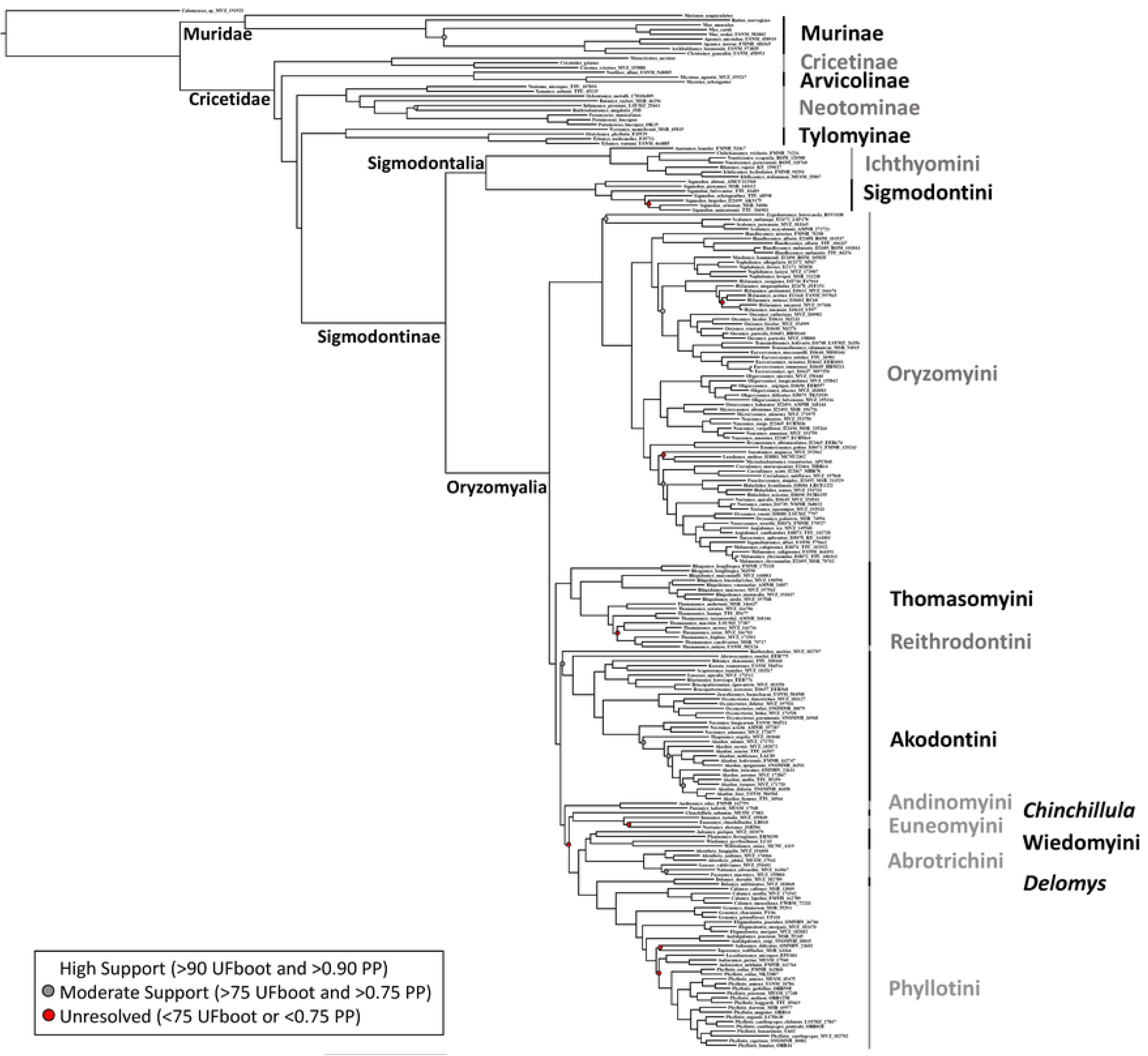
Phylogeny derived from all AHE loci with branch lengths generated by IQ-tree. Nodes with greater than 90% ultrafast bootstrap support (UFBoot) in IQ-tree and greater than 0.90 local posterior probably (LPP) in Astral-III are represented by the absence of dots on the nodes, all nodes with between 75 - 89% UFBoot and 0.75 – 0.89 LPP are represented with a gray dot, and all nodes with less than 75% UFBoot and / or 0.75 LPP are represented with a red dot. Taxonomic groups are labeled including all subfamilies of Cricetidae and all tribes of Sigmodontinae.

**Fig 2:**
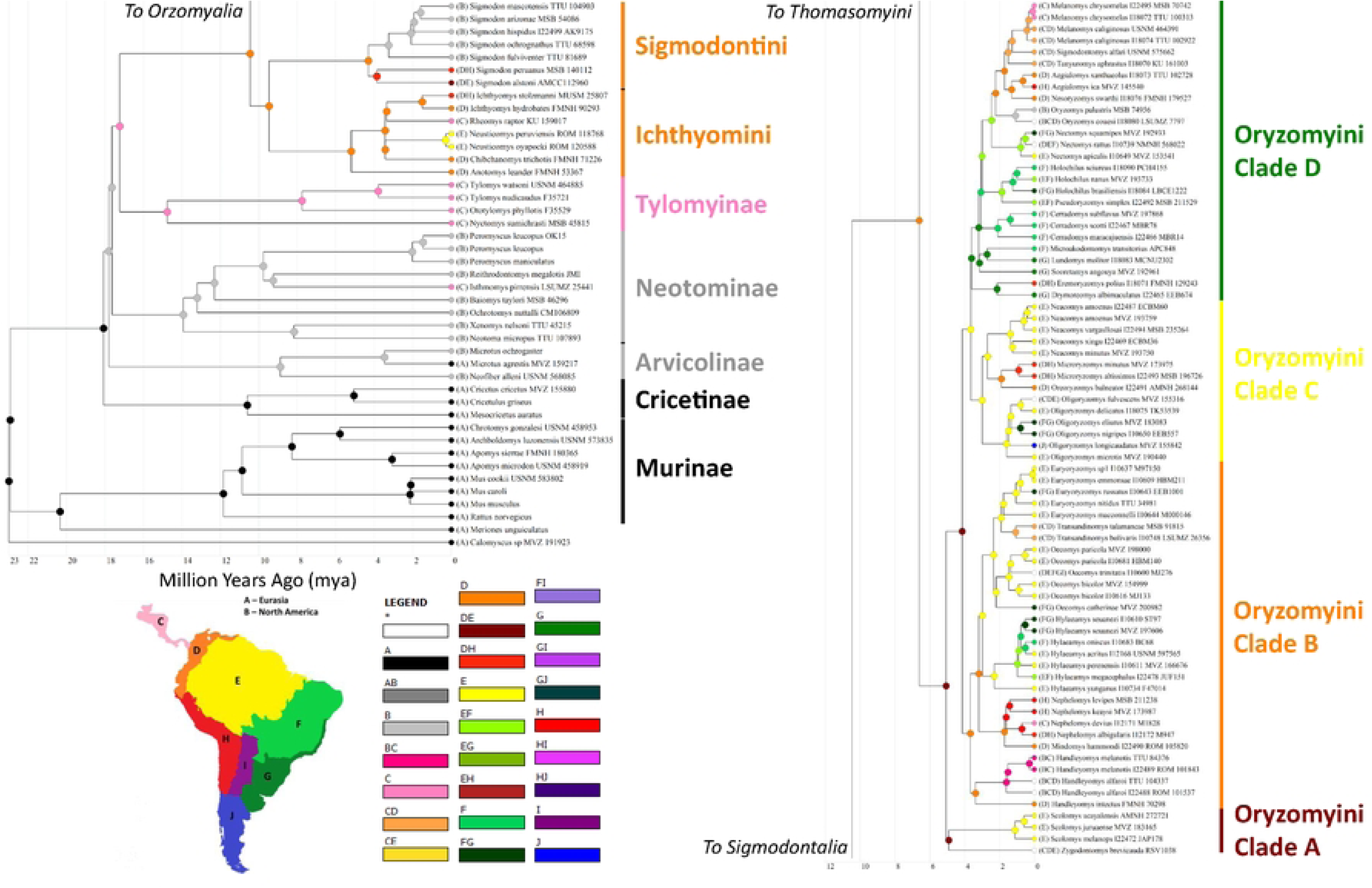
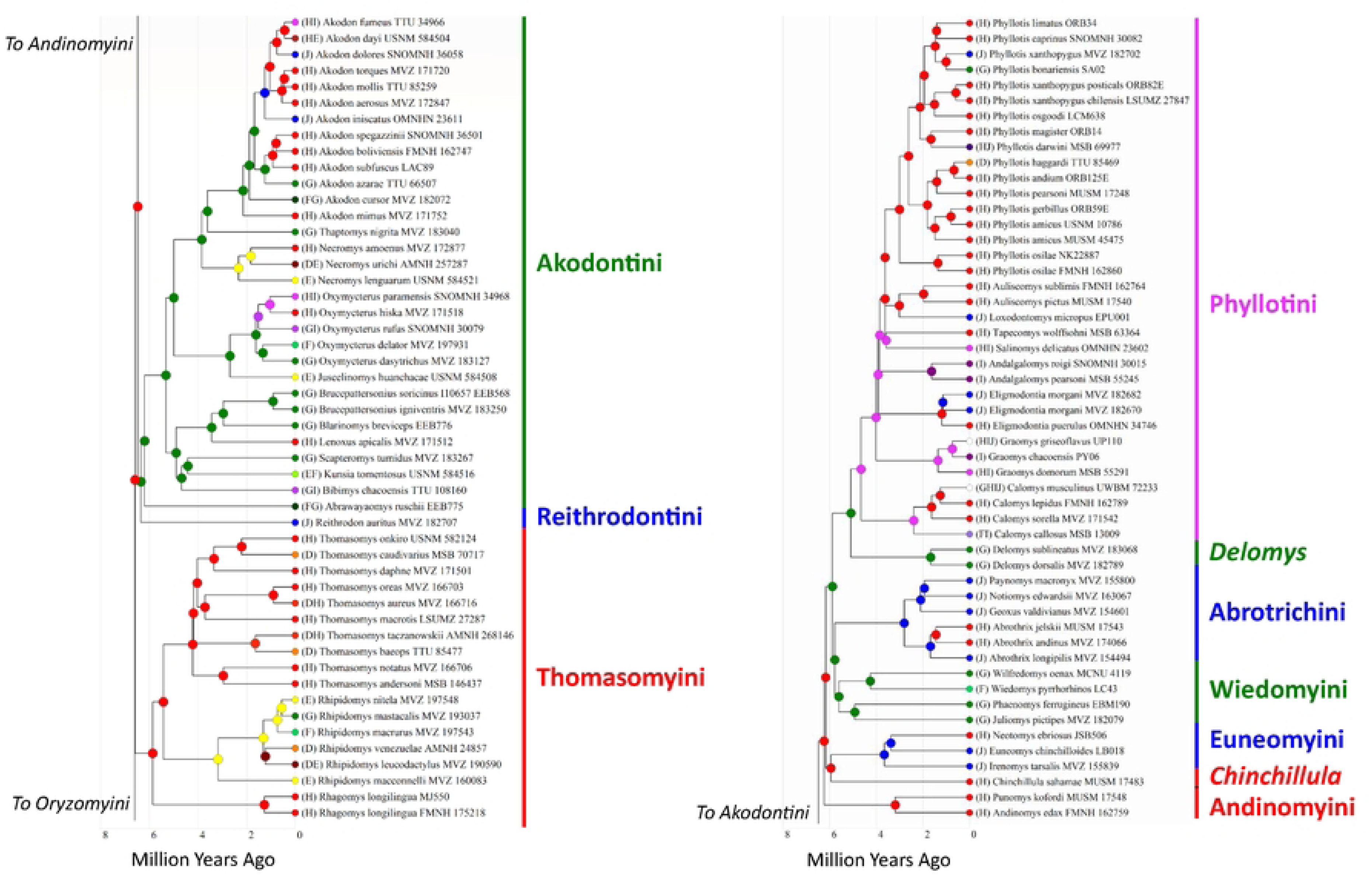
Fossil calibrated maximum likelihood phylogeny generated with ancestral biogeographic reconstruction. Circles on nodes represent highest probability reconstruction of ancestral range estimated using a full hierarchical Bayesian Binary MCMC (BBM) model in RASP. Branches colored to match the map of coded regions. Taxonomic groups (tribes and subfamilies) are labeled and colored based on the MRCA of the group. Oryzomyini is split into the four clades as described in Percequillo et al. [41] given the size of the tribe and the divergent ranges of the clades within the tribe.

Regarding the three *incertae sedis* genera (*Abrawayaomys*, *Chinchillula*, *Delomys*), all three were sister to established tribes, with *Abrawayaomys* sister to Akodontini, *Chinchillula* sister to Euneomyini, and *Delomys* sister to Phyllotini (Fig 1).

### Fossil calibrated phylogeny

The crown group of Cricetidae dated to 18.08 mya (15.11 – 21.64 mya 95% CI) with all five subfamilies diverging by 17.24 mya (13.17 – 21.64 mya 95% CI). Sigmodontinae and predominantly Central American Tylomyinae were the last to diverge (Fig 2). Within Sigmodontinae, the first split, between Oryzomyalia and Sigmodontalia, was recovered at 10.46 mya (8.25 – 13.27 mya 95% CI) with Oryzomyalia radiating nearly four million years later during the Late Miocene at 6.61 mya (5.49 – 7.96 mya 95% CI). The first split in Sigmodontalia was earlier than Oryzomyalia at 9.47 mya (7.33 – 10.46 mya 95% CI), with the tribes Sigmodontini and Ichthyomyini diversifying well after (at 4.31 mya and 5.20 mya, respectively). The initial radiation in Oryzomyalia generated all 10 remaining tribes within ∼825,000 years during the late Miocene (represented by gray numbered circles; Fig 3) except for Phyllotini, which split from *Delomys* around 5.1 mya (3.96 – 6.59 mya 95% CI, gray circle seven; Fig 3).

**Fig 3:**
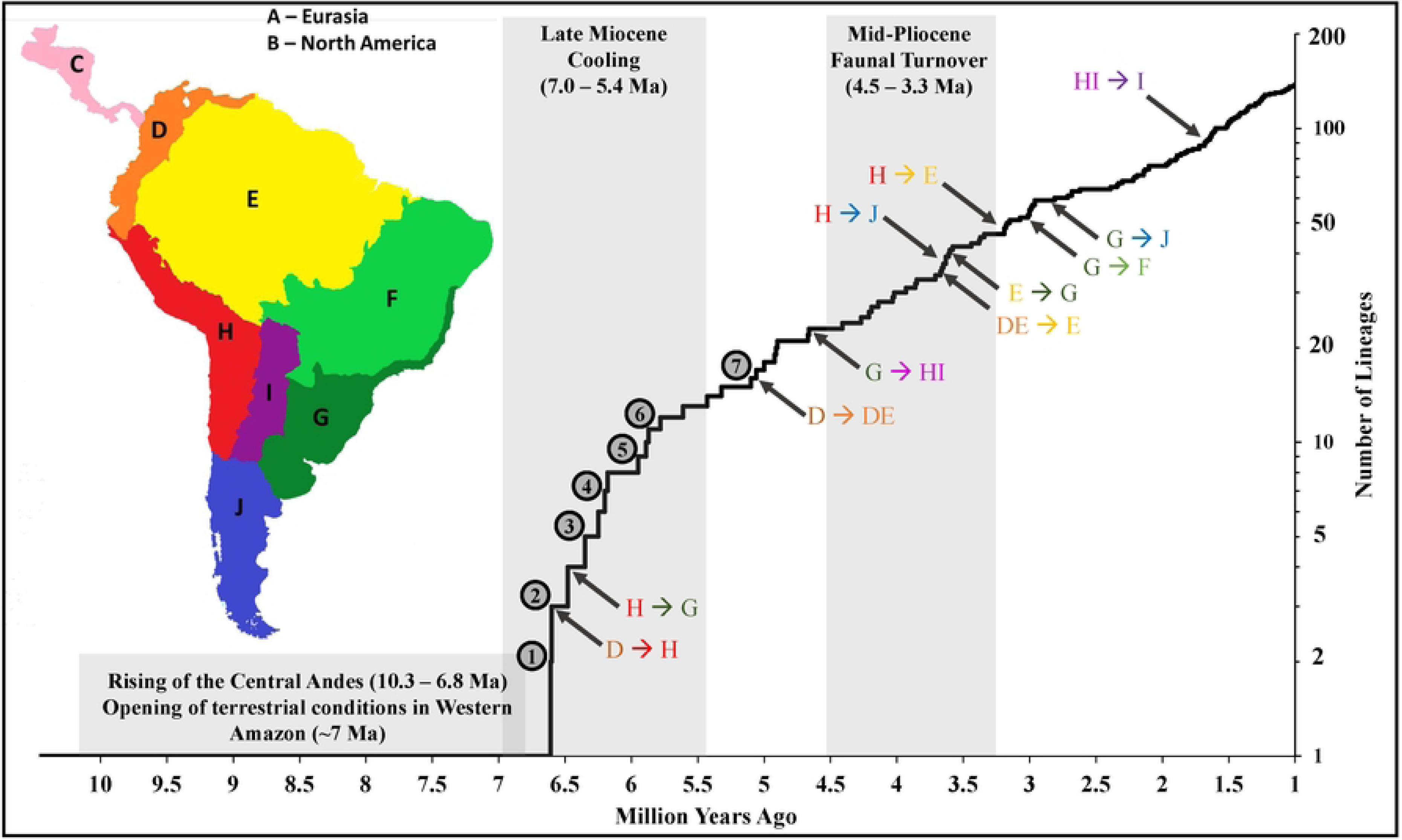
Lineage through time plot of Oryzomyalia starting at the origin of the clade (10.46 mya) to 1.0 mya before present. Origin of the crown groups for tribal-level clades are labeled with numbered circles (1 = Oryzomyini, 2 = Thomasomyini, 3 = Reithrodontini and Akodontini, 4 = Andinomyini, 5 = Euneomyini and Chinchillulini, 6 = Abrotrichini and Wiedomyini, and 7 = Delomyini and Phyllotini). First transitions into new regions in the ancestral biogeographic reconstruction (Fig 2) are labeled on the plot with gray arrows; map of regions included for reference. Time periods for geological and biological events discussed in the text are indicated by gray background.

### Ancestral biogeographic reconstruction

The earliest node to be reconstructed as South American was the MRCA of Sigmodontinae around 10.46 mya in the northernmost region (D), followed by Sigmodontalia at 9.47 mya, and well before the later radiation of Oryzomyalia at 6.61 mya (Fig 2). The initial radiation of Oryzomyalia saw the first split between Oryzomyini, in the northernmost region (D), and the rest of Oryzomyalia spreading down the Central Andes (H; Figs 2 and 4A). This later group then crossed into the temperate Atlantic Forest and grasslands (G) twice (Figs 2 and 4A); once by the Reithrodontini-Akodontini clade about 6.35 mya (4.95 – 7.96 mya 95% CI) and then by the Wiedomyini-Abrotrichini-*Delomys*-Phyllotini clade around 5.89 mya (4.35 – 7.96 mya 95% CI). Over the next ∼1.1 mya (4.67 – 5.77) no transitions between regions occurred, with new lineages remaining in the same region as their ancestors, apart from Oryzomyini which expanded into the Amazon and Guianan tropical forest (E) during this period (Fig 4B). This was followed by a period in the Mid-Pliocene where all remaining regions of South America were colonized, starting around 4.66 mya with the spread of Phyllotini into the dry regions of central South America (HI) and ending with the colonization of the Cerrado (F) by Oryzomyini 3.07 mya (Fig 4C). This period also included the first colonization of Sigmodontinae back into Central and North America by the cotton rat *Sigmodon* around 4.31 mya (3.24 – 5.73 mya 95% CI), a genus who’s 14 species primarily occur today in North America, as well as the fish-eating rat *Rheomys* around 3.44 mya (2.48 – 4.76 mya 95% CI, a genus of four Central American species. After this period, diversification within and across regions of South America occurred continuously with several taxa moving into Central America, namely species in genera *Melanomys*, *Nephelomys*, *Oligoryzomys*, *Sigmodontomys*, *Tanyuromys*, *Transandinomys*, *Zygodontomys* (all being members of Oryzomyini), and *Akodon*, and two additional transitions further into North America by Oryzomyini (*Handleyomys* and *Oryzomys*; Fig 2).

**Fig 4:**
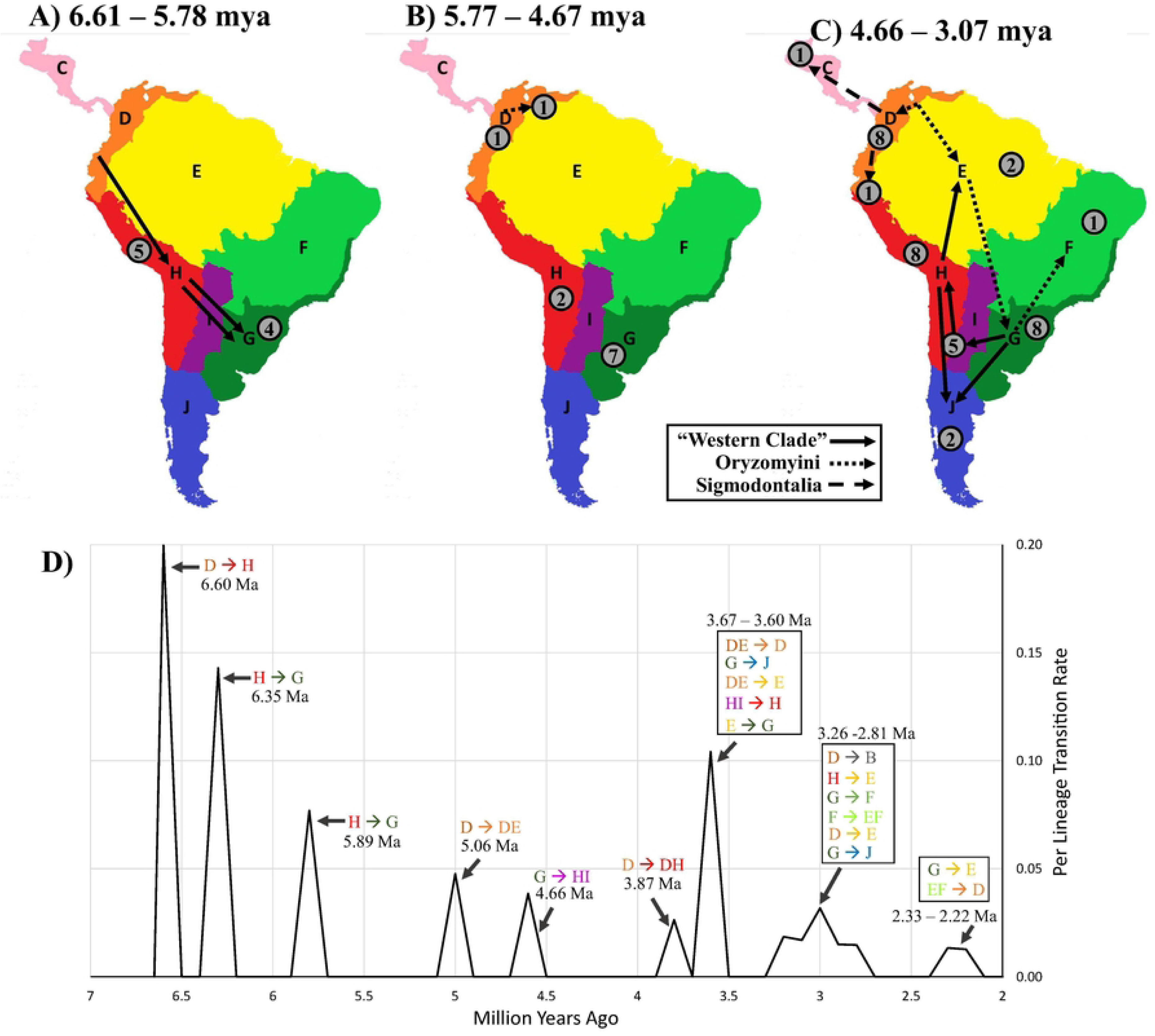
Maps of Sigmodontinae radiation over time. Figures start at the basal split in Oryzomyalia (6.61 mya) to the point when the last biogeographic region was colonized in the ancestral biogeographic reconstruction (3.07 mya; Fig 2) as well as the lineage transition rate (number of transitions per branch at the start of the time frame per million years recorded in 100,000-year intervals; Fig 4D). Maps are split into three time periods; (4A) initial Late Miocene radiation (6.61 - 5.78 mya), (4B) period of low among-region dispersal (5.77 – 4.67 mya), and (4C) mid-Pliocene colonization (4.66 – 3.07 mya). Arrows on the maps represent all transitions between regions in the biogeographic reconstruction (e.g., each represents a branch in the reconstruction that has a node before and after that differ in region) and are split into three groups: Sigmodontalia (dashed arrow), Oryzomyini (dotted arrow), and the rest of Oryzomyalia (solid arrow, “Western Clade”). Gray circle with numbers indicates the number of new lineages that originated within the region during the time frame (e.g., *in situ* splits in the phylogeny, which are the generation of new branches that have nodes reconstructing to the same region before and after). All circles are placed within a single region with the exception of a circle in the top of the center map for D/E (1 speciation event) with arrows splitting out of the combined region in the right most map and a circle for H/I (5 speciation events) in the last map. An arrow from G to J is included in the last map because of the presence of the first fossils of mono-generic Reithrodontini during this time period even though the reconstruction in Fig. 2 does not explicitly indicate dispersal during this period.

## Discussion

Sigmodontinae represents the largest subfamily of Neotropical mammals and the largest clade of mammals to participate in the Great American Biotic Interchange [6, 58], an event that shaped communities throughout the Americas to their modern composition. The subfamily represents one of the most recent continental scale radiations in mammals and provides an opportunity to understand the processes that modelled through time and space a continental radiation, a rarity compared to island radiations. While in the recent past, reconstructing continental radiations was challenging given the scale and complexity of these radiation, the advancements of next generation sequencing and paleoclimatic reconstructions, allied to adequate samples in museums and collections, provide an opportunity to finally disentangle these complexities and reconstruct challenging histories.

Here we present a highly supported genomic phylogeny using > 83% of all genera and ∼ 40% of the > 500 species described to date, resolving the relationships among tribes, placing all *incertae sedis* taxa, and reconstructing the sequence of dispersals and lineage generation across South America. Below we discuss what this reconstruction means in light to prior phylogenetic work and the changing landscape of South America in order to understand potential role of changing climatic, biologic, and geologic conditions.

### Comparison to prior phylogenies

The radiation of Sigmodontinae has been challenging due to short internode distances at the base of the radiation, resulting in 14 different reconstructions of the tribes at the base of Oryzomyalia (the clade responsible for generating most of the diversity in the South American radiation) since 2012 [9–11, 13, 17, 18, 27, 28, 30, 31, 34, 35]. Parada et al. [13] showed that the use of next generation sequencing can resolve these nodes, producing the first high support for the relationships between the tribes of Oryzomyalia. While their taxonomic sampling (53 species in 36 genera) was sufficient to clarify some of the most important phylogenetic questions, some geographically critical taxa were missing and therefore precluded an investigation of changes in diversification and transitions over landscapes. Vallejos-Garrido et al. [18] were able to include many of the critical missing species in Sigmodontalia by combining the Parada et al. [13] data set with other published single-gene sequences from Sanger sequencing. In contrast, the phylogeny generated here, while congruent with Parada et al. [13], captures nearly all early nodes in the radiation (e.g., Miocene and Pliocene), and with many more genes for most remaining species than in Vallejos-Garrido et al. [18]. While there are still many species missing in this phylogeny, nearly all of these represent branches within genera that originated in the last two million years. Thus, this phylogeny allows for a comprehensive evaluation of the timing and transitions of the South American radiation throughout much of the early phases of lineage generation discussed below.

This phylogeny also provides a framework for an update to the systematics of the subfamily that was not previously possible and while a thorough discussion of all updates to the systematics is outside the scope of this work, we do feel a discussion of major changes in tribal systematics is warranted, particularly in regard to *incertae sedis* taxa that have not previously been sampled in prior molecular phylogenies. Thus, we provide a revised classification along with a discussion of these decisions.

### Migration into South America

The subfamily Sigmodontinae belongs to a larger clade of New World mice and rats, that also includes the largely North American subfamily Neotominae and the largely Central American subfamily Tylomyinae, the latter to which the predominantly South American Sigmodontinae is sister (Fig 1). While most species in Sigmodontinae occur in South America, with the largest clade (Oryzomyalia) having a well-established South American origin, the subfamily as a whole has been often thought of having a Central or North American origin due to the earliest apparent sigmodontine fossils being located in North America [16, 21, 59–61] or due to the presence of the remaining tribes, grouped as Sigmodontalia, having an hypothesized Central or North American origin [11]. Steppan et al. [7] suggested that three parallel colonization events into South America occurred, with one being the cause of the rapid radiation in Oryzomyalia, most similar to the earlier model of Marshall [62]. In contrast, our reconstruction suggests the more parsimonious explanation of a single South American invasion by Sigmodontinae, with the crown group reconstructed in northern South America during the late Miocene (Fig 2). This change in reconstruction is due to the recovery of Sigmodontalia also as South American, sister to a previously recognized South American radiation of Oryzomyalia. The driving factor for the difference in reconstructed distributions is that this is the first subfamilial study to include all basal branches of Sigmodontalia, including previously unsampled South American Ichthyomyini and South American forms of the genus *Sigmodon* (but also see Salazar-Bravo et al. [63] who thoroughly sampled Ichthyomyini). Most previous studies across Sigmodontinae have included only two or three genera of Sigmodontalia and fewer than five species with a bias of sampling North and Central American species. This bias is an issue in Ichthyomyini where 13 of the 17 species occur in South America yet most studies have utilized the only Central American lineages (e.g., [8, 9, 11, 34]). Although Vallejos-Garrido et al. [18] included South American species of *Sigmodon*, they lacked the critical South American Ichthyomyini species, and as a result their reconstruction of the MRCA of Sigmodontinae was uncertain regarding continent or origin.

An important rationale for a Central or North American origin of Sigmodontinae is that the oldest putative fossil, *Prosigmodon*, occurred in Mexico [21, 61]. However, this fossil is known from sediments with ages well after the origin of the subfamily, with the original description [61] referring to it as being from Late Hemphillian to Early Blancan, but without specific dates. A later review [64] placed these earliest fossils in the “Latest Hemphillian” strata, around 5.3 – 4.9 mya. Our reconstruction can explain this fossil, since the presumed dates fall within the 95% CI estimate of the transition of *Sigmodon* into North America (5.73 – 3.24 mya); thus, the presence of this fossil does not conflict with a South American origin. The lack of Miocene fossils in Sigmodontinae also fits with our reconstruction, since the subfamily would not have expanded into regions with adequate fossil bed formations (e.g., North America and southern South America) until the early Pliocene, around the time we first see fossils in both locations [65–68].

A single South American transition at the crown of Sigmodontinae challenges the long-standing notion of invasion as a trigger for the radiation because our reconstruction indicates a nearly four-million-year gap between the colonization of South America (10.46 mya; 8.25 – 13.27 mya 95% CI) and the radiation of Oryzomyalia (6.61 mya; 5.49 – 7.96 mya 95% CI). If not due to recent invasion of a new continent, why was there a dramatic increase in lineage creation and transition rate in Sigmodontinae in the Late Miocene (e.g., what triggered the Oryzomyalia radiation)? A less parsimonious reading of the reconstruction could be that as implied by Marshall [62] and favored by Steppan et al. [7, 8], there were several expansions into South America in parallel (although not necessarily simultaneously) followed by local extinction in Central America. This scenario is less likely if the three events were entirely independent, but more likely if they shared an underlying mechanism of propensity to disperse via island hoping combined with possible habitat change shrinking the northern distributional limits.

### What sparked the Oryzomyalia radiation?

Based on our reconstruction, Oryzomyalia began radiating around 6.61 mya (95% CI 7.96 – 5.49 mya), going from a single lineage to 12 in less than one million years (∼850,000 years; 6.61 – 5.78 mya), resulting in most of the tribal diversity for the group (Fig 2). This radiation corresponds with Late Miocene Cooling (7.0 – 5.4 mya) and the collapse of the Pebas mega-wetland system (∼6.8 mya) in Amazonia [37]. Recent climate simulations [36], paleo-pollen surveys [69], and sediment analyses [37] suggest a decrease in aquatic habitats in northern South America (e.g., wetland and marshes replaced with forest and grasslands) and spread of montane and pre-montane forest in the Northern Andes resulting in a connection with those in the Central Andes [37]. The Pebas mega-wetland separated the northern proto-Andes from Amazonia and the Guianan region, potentially restricting early sigmodontines to a north-western peninsula. Consequently, the ecological opportunity presented by the continent of South America (sensu Schenk et al. [8]) may not have been accessed until the collapse of the Pebas system, delaying the burst of diversification from the initial colonization. In general, while there was an increase in aridity in South America during the Late Miocene Cooling, resulting in spread of grasslands especially in the northern lowlands and the southern latitudes [36], regions where the clade radiated correspond to areas of increased precipitation (orographic rains brought by SE trade winds, based on deposits on the slopes of Serra do Mar in along the southern coast of Brazil and the southern Andes) and forest expansion [37]. This suggests that these early temperate, tropical and subtropical forests of the region “G” (Fig 2) and of the montane and pre-montane forests on the eastern slopes of the Central Andes (“H” Fig 2) played a key role in early diversification of the group.

Our estimate for the arrival of sigmodontines into the more temperate habitats of region “G” around 6.35 mya is consistent with a revised time scale of the fossil record for this region. Prevosti et al. [70] date the earliest South American cricetid fossil (unidentified Sigmodontinae) from the Cerro Azul Formation (upper Huayquerian) of Buenos Aires Province to 6.935–5.464 mya (median age 6.192 mya). This still leaves the often-noted absence of fossils during the 4 My gap between first colonization implied by molecular estimates and the earliest South American fossils, but if the early lineages were associated with lowland forests and cloud forests along the Andes (region “D”), then an absence of fossils is not wholly unexpected.

Alternatively, the spread of grasslands during Late Miocene could have played a role in the initial radiation; however, this does not explain the more than three-million-year lag between first colonization in the north and the reconstructed dispersal into to the southern regions (e.g., region “G”), which were dominated by grasslands at the time [36] as they still are today. Instead, the reconstruction suggests the existence of a corridor of montane and temperature forest connecting the Central Andes to southeastern portion of South America during the Late Miocene, leading to the origin and diversification of most of the initial tribes in Oryzomyalia, with a later colonization of the rest of the continent during the Mid-Pliocene.

Similar timing of diversification has also been observed in the South American fauna and flora, particularly those endemic to cloud forests. Recently, phylogenetic studies of taxa associated with cloud forests have found that the late-Miocene was an important time for early radiations across the continent, connecting the mid-montane Brazilian Highlands (in region “G”) with that of the North and Central Andeans (regions “D” and “H” respectively). This included cloud forest endemic palms (*Geonoma* [71]), hummingbirds (*Anelomyia*, [72]), and other flowering plants (e.g., *Macrocarpaea*, [73]). The gentian genus *Macrocarpaea* shows the most striking similarity to the timing and location of the early radiation in Oryzomyalia, but is reverse, going from a single lineage in the Brazilian Highlands to the Central Andes and then radiating into five new lineages in a period of ∼1 million years starting ∼7.2 mya. However, unlike Sigmodontinae, these species remain endemic to South and Central American cloud forest.

The importance of cloud forest early in the evolution of Sigmodontinae can also be seen in the sister subfamily Tylomyinae that is largely endemic to the cloud forest remnants of Central America. Based on paleopollen and phylogenetic studies, these Mesoamerican cloud forests were present in Central American as early as 20 mya and far more abundant across Central American in the Miocene and Pliocene [74–76]. Thus, if we look at the paleoclimatic conditions during the timing of the split of Sigmodontinae and Tylomyinae in Central America around 17.24 mya (13.17 – 21.64 mya 95% CI), we see that cloud forests were abundant in the landscape of Central America as well as emergent in Northern South America, but cut off from the rest of South America until the late Miocene uplift of the Central Andes. That coincided with cooling and increased precipitation of the period leading to range expansion of cloud forest that later receded in the Pliocene and Pleistocene [36, 37, 77]. Thus, our reconstruction coupled with paleoclimatic reconstructions of Miocene and Pliocene Central and South America, as well as similar timing of phylogenetic radiations of cloud forest endemics, suggest that cloud forests played a prominent role in the radiation of Sigmodontinae, an idea that has been obscured by the age and complexity of the radiation as well as the continental range of Sigmodontinae observed today. This leads to the question, what sparked the spread of this group throughout South American in the latter Pliocene in contrast to their Central American relatives that remained largely endemic to cloud forest as they receded in the latter ages?

### Mid-Pliocene diversification

At the start of the Pliocene (5.33 mya), most of the tribal lineages of Sigmodontinae had arisen but were constrained to either northern South America (“D” / “DE”; Sigmodontalia and Oryzomyini), the Central Andes (“H”; Thomasomyini, *Chinchillula*, Euneomyini, and Andinomyini), or Southeastern Atlantic (“G”, Akodontini, Reithrodontini, Wiedomyini, Abrotrichini, and *Delomys* + Phyllotini), presumably largely in cool moist forests that dominated these regions at the time. It was not until the Mid-Pliocene (4.66 – 3.07 mya) that sigmodontines spread across all the remaining regions of the continent, including nearly all of the diverse South American ecosystems harboring these rodents today, from lowland deserts to the highest elevation for any mammal, and from lush rain forest to vast grasslands. This expansion often occurred with a single lineage spreading into a new region followed by small bursts of diversification in these groups throughout this time frame (4.66 – 3.07 mya). For example, Phyllotini spread to the open semi-arid and arid areas of the Dry Chaco and Central Andes (“H/I”) 4.66 mya (95% CI 5.97 – 3.94 mya) and radiated from one to eight lineages over the next million years (Fig 2). Similarly, Oryzomyini had several radiations after spreading into new regions, e.g., the origin of “Clade D” around 3.60 mya (95% CI 4.69 – 2.77 mya) after a spread into the Southeastern Atlantic regions (“G” and “F”) resulting in six lineages from a single lineage in ∼600,000 years (Fig 2).

These bursts resulted in an increase in both lineage diversification rate, as measured by the number of new lineages per branch per million years, and lineage transition rate, as measured by the number of new regions occupied per branch per million years (Fig. 4C), during the Mid-Pliocene. For example, the average lineage diversification rate (0.91 new lineages per branch per million years) during this time (4.66 – 3.07 mya) was higher than the period between the initial radiation (5.77 – 4.67 mya; 0.71 new lineages per branch per million years) and after the Mid-Pliocene burst (3.06 – 2.00 mya; 0.44 new lineages per branch per million years). Similarly, the transition rate was more than double during this time compared to the periods before and after (0.23 versus 0.06 and 0.10 respectively). While neither of these metrics were as high as during the initial Miocene radiation (6.61-5.78; 7.23 new lineages per branch per million years and 1.81 transition per lineage per branch per million years), the Mid-Pliocene still reveals a second set of radiation and expansion events. (Fig 4). Schenk and Steppan [11] identified the correlation of these two aspects, with a strong peak at the base of Oryzomyalia, but with less clear support for the mid-Pliocene peak discovered here. Results from Vallejos-Garrido et al. [18] also indicate increased among-region dispersal rates during this same period for most of their biogeographic regions. We suspect the greater distinctness of the second set of bursts seen here is a consequence of improved estimates of relative ages for nodes and a more accurate geographic reconstruction because of the different and better supported topology. Nearly every biogeographic reconstruction in recent years has also employed different regionalization schemes, making it more difficult to compare results, as was noted by Vallejos-Garrido et al. [18].

During the Mid-Pliocene (4.66 – 3.07 mya), sigmodontines spread throughout the continent with the Oryzomyini spreading through the eastern regions and the rest of Oryzomyalia spreading through the southern and western regions (Fig 4C). This time period coincided with high extinction of South American mammals in a period known as the Mid-Pliocene Faunal Turnover in which more than half of all mammalian species became extinct (4.5 mya – 3.3 mya [78]). Shifting climates throughout the continent resulted in changes in seasonal rainfall in the Central Andes and Amazon. The consequence was a further decline of wetland habitats [38] and increasing aridity in the southern regions (particularity “F”, “I”, “J”, and southern and eastern parts of “H”) resulting in expanding grasslands [36]. The decline of mammalian competitors and predators coupled with muroid rodents’ high reproductive rate and ability to survive in these new habitats likely contributed to this Mid-Pliocene burst and the continental range of the group today. This insight changes the narrative from one of initial burst and rapid spread throughout the continent to one of episodic expansions and radiations, taking advantage of changing climate, flora, and fauna.

### Systematics of tribes and placement of *incertae sedis* lineages

Currently, twelve tribes are generally recognized in the subfamily Sigmodontinae, namely Sigmodontini (Wagner 1843)[79], Akodontini, Ichthyomyini (Cockerell and Printz 1914)[80], Oryzomyini, Phyllotini (Vorontsov 1959)[81], Thomasomyini (Steadman and Ray 1982)[82], Wiedomyini (Reig 1980)[20], Abrotrichini (D’Elía *et al*. 2007), Euneomyini (Pardiñas *et al*. 2015)[83], Andinomyini (Salazar-Bravo *et al.* 2016)[84], Reithrodontini (Cazzaniga *et al.* 2019)[85], and Neomicroxini (Pardiñas *et al*. 2021)[39]. All polytypic tribes were recovered here as monophyletic. The monophyly of these tribes have similarly been confirmed with similar large taxon sampling of small multi-gene phylogenies [11] as well as recent genomic studies employing smaller taxonomic sampling [13, 41] and a combined-data analysis [18]. However, three genera (*Chinchillula*, *Abrawayaomys*, and *Delomys*) remained *incertae sedis* in past studies due to not being sampled in the multi-locus studies or low support values for the placement of the genera. Below we discuss the assignment of the three *incertae sedis* taxa and provide brief justification of updates in their tribal placement in order to resolve the tribal systematics of the subfamily.

The Linnean rank of a clade is an arbitrary decision as long as doing so conforms to several rules and guidelines (e.g., [86]). Such decisions then are best evaluated on the information conveyed by the rank and the utility to systematists. The decision often is a tradeoff between providing grouping information (lumping) versus recognizing distinctiveness, whether phenotypic or temporal (splitting). More inclusive taxa contain more variation, and therefore can be more difficult to define phenotypically, eroding the phenotypic information communicated. Less inclusive taxa necessarily contain less grouping information, and a proliferation of names can devalue the utility of each. In addition, some clades are of such scientific interest that they are frequently referred to, and having a name simplifies communication. As an example, the rank-less clade Oryzomyalia was named by Steppan et al. [7] because it contained a large proportion of sigmodontine diversity, was separated from its sister group by a relatively long period of independence, and the base of which involved a rapid diversification, making it of particular scientific interest, but for which no succinct morphological diagnosis was available. Providing a name has proved useful to communication. In the following recommendations and accompanying classification for the subfamily, we strive to maximize grouping information, avoid taxonomic inflation, retain common usage where possible to maintain stability, but still recognize distinctness. Pardiñas et al. [87] recommended erecting several tribes for single but morphologically distinctive genera. In some cases, we agree, in others we concluded that distinctness was overemphasized and thus information was eroded, but in no case were any of their recommendations incorrect.

All three *incertae sedis* genera were recovered outside of established tribes, with *Chinchillula* sister to Euneomyini, *Abrawayaomys* sister to Akodontini, and *Delomys* sister to Phyllotini. In all cases the support for these arrangements was high (> 95% bootstrap support and 0.95 LPP), providing confidence in their placement. Thus, a discussion of whether to expand the breadth of the established tribes or providing new tribal placement is warranted.

### Genus *Chinchillula*

*Chinchillula* and members of Euneomyini, to which it is sister, share many morphological characteristics, such as dense and lax pelage, long incisive foramina, and simplified terraced molars. However, many of these characteristics are shared with other Andean species (e.g., Andinomyini and Phyllotini), likely convergent traits to herbivory and high montane open habitats, that previously resulted in their placement within the more inclusive concept of the tribe “Phyllotini” (e.g., [20, 22, 88–90]), that is now known to be polyphyletic [7, 83, 84, 91]. Recent molecular reevaluation on the contents of the tribe Phyllotini based on few genes [83, 84], resulted in the description of two new tribes, namely Euneomyini and Andinomyini, that are supported by the shared molar complexity within each of these groups, as well as the redefinition of a less inclusive concept of the tribe Phyllotini ([83, 84]; genomic data here presented supported this decision). *Chinchillula* presents a unique color coat pattern within sigmodontines, as well as highly specialized molar complexity that, if anything, resembles that of Andinomyini rather than Euneomyini [92], in contrast to the molecular phylogeny. Steppan [22] regarded *Chinchillula* as a Phyllotini *sedis mutabilis*, emphasizing the uncertain affiliation of this genus to *Auliscomys* (presently deeply nested within Phyllotini) and *Andinomys* (member of tribe Andinomyini). The unique morphology, coupled with the early split from Euneomyini (5.95 mya) and long branch separating them from Euneomyini, leads us to suggest a separate tribe consisting of a single species *Chinchillula sahamae*.

> Tribe Chinchillulini, new tribe

> Type genus: *Chinchillula* Thomas, 1898

> Contents: *Chinchillula* Thomas, 1898

Definition: this new tribe of subfamily Sigmodontinae is a member of a clade containing the tribes Andinomyini, Euneomyini, Wiedomyini, Abrotrichini, *Delomys* and Phyllotini. Chinchillulini is sister to the tribe Euneomyini, a relationship with high support on the genomic analysis here presented.

Diagnosis: as a monotypic tribe, Chinchillulini is diagnosed by the unique combination of traits for the genus, as follows: unique pelage color pattern, with distinct white patches on the muzzle, neck, anterior surface of legs and rump, that are prolongations of the predominantly white ventral region; dark band-shaped patches present on the rump; pelage very long, dense and lax; very long pinnae, with distinct tuft of white hairs on its basal portion; mystacial vibrissae very long and dense; superciliary and genal vibrissae present; eye ring fine and dark; manus and pes densely covered by white/silvery hairs, digits with long and dense ungueal tufts; tail much shorter than head and body length (30-50%), with a light brown stripe on the dorsal portion; skull very robust, with broad rostrum, not inflated frontal sinuses; nasals broad tapering posteriorly; interorbital region long and narrow, convergent posteriorly, with squared or rounded supraorbital margins; interfrontal fontanelles absent; fronto-parietal suture V-shaped, narrow; interparietal robust, trapezoidal; alisphenoid strut variably present, buccinator-masticatory foramen and accessory oval foramen separated or confluent; anterior opening of the alisphenoid canal open; stapedial foramen, squamoso-alisphenoid groove and sphenofrontal foramen present; posterior opening of alisphenoid canal large and conspicuous, with deep posterior groove or depression; secondary anastomosis of internal carotid artery not visible on dorsal surface of parapterygoid plate; parapterygoid fossa deep; incisive foramina very wide and very long extending to the anterocones of M1; mesopterygoid fossa wide, equal or narrower to parapterygoid fossae; mesopterygoid fossa extending anteriorly to maxillaries, but not reaching alveolus of molars; anterior margin squared; palate without deep groves or fossa; mandible with long and narrow condylar process; upper incisors very broad and quite robust, with distinctive orange enamel band; orthodont; maxillary toothrows parallel or slightly convergent anteriorly; labial and lingual flexi of upper molars not penetrating at molar midline; molar series with well-developed coronal hypsodonty and plane occlusal surface; lingual cusps in subadults slightly anterior to the labial cusps; molars trilophodont, very simplified, with very deep and wide labial and lingual flexi, without labial and lingual cingula or styles, mandible with long and narrow condylar process, vertebral formula 13 thoracic, 6 lumbar and between 24-29 caudal vertebrae (see [22, 33, 89], for additional traits).

Known collection localities: Pacheco et al. [93] identified nearly 50 localities in Bolivia, Chile and Peru, from Junín Department in the north to Tarapaca in the south, and ranging in elevation from 3,550 m to 5,100 m. Salazar-Bravo [94] noted that Hershkovitz [89] mapped the locality of Sumbay, Arequipa, at about 2,000 m, likely an error, as there are not known records of this species below 3,550 m ([95] indicates ca. 4,000 m as the elevation of Sumbay). A closer inspection of Hershkovitz’s ([89]: fig. 117) map shows that localities (3) Arequipa and (4) Sumbay, are not only misplaced, but should be reversed. Additionally, the lowest elevation from Cordillera de Sicuani, Cusco, at 3550 m, is almost 400 m of elevation lower than all known records [93] and should be confirmed.

### Genus *Abrawayaomys*

*Abrawayaomys* has been difficult to place given limited voucher material and the fact that it shares many characteristics with thomasomyines, particularity with *Rhagomys* (especially on molar topography, as discussed by Percequillo et al. [96]) as well as several cranial characteristics similar to those exhibited by the members of tribe Akodontini (see Pardiñas et al. [97] and Percequillo et al. [96]). This has led to the tentative placement with Thomasomyini, but strong convergence with Akodontini was assumed due to similar diets and ranges [97]. As pointed out by Percequillo et al. [96], both morphological and molecular phylogenies have been conflicting and with low support, leading to uncertainty as to which characters represent synapomorphies versus homoplasy. Here we find strong support for the sister-taxon relationship of *Abrawayaomys* with Akodontini and argue that the shared characters of “inflated frontal sinuses, hour glass shaped interorbital region with a broad interorbital constriction, U-shaped coronal suture, and some traits of the simplified molar occlusal pattern” ([97]: pg. 58), as well as the large postglenoid foramen and subsquamosal fenestra, small posterolateral palatal pits and the tegmen tympani overlapped to the posterior suspensory process of squamosal, recognized by Percequillo et al. [96], represent synapomorphies. Therefore, we suggest the inclusion of the genus *Abrawayaomys* within the concept of tribe Akodontini, based on the genomic evidence, shared morphological features, and overlap in biogeography.

> Tribe Akodontini

New contents: *Abrawayaomys* Cunha and Cruz, 1979; *Akodon* Meyen, 1833; *Bibimys* Massoia, 1979; *Blarinomys* Thomas, 1896; *Brucepattersonius* Hershkovitz, 1998; *Deltamys* Thomas, 1917; *Gyldenstolpia*, Pardiñas et al., 2009; *Juscelinomys* Moojen, 1965; *Kunsia* Hershkovitz, 1966; *Lenoxus* Thomas, 1909; *Necromys* Ameghino, 1889; *Oxymycterus* Waterhouse, 1837; *Podoxymys* Anthony, 1929; *Scapteromys* Watherhoude, 1837; *Thalpomys* Thomas, 1916; *Thaptomys* Thomas, 1916.

### Genus *Delomys*

*Delomys* was monophyletic and represents the most recent divergence of any of the *incertae sedis* taxa, splitting from Phyllotini around 5.1 mya. *Delomys* split from Phyllotini before the latter radiated into the open and often arid areas of South America at the start of the Mid-Pliocene Faunal Turnover. This left *Delomys* in the Brazilian Atlantic Forest while phyllotines adapted to these new open and arid environments, developing morphological traits related to these new environments, some of which were unique to the tribe while others converged with other Andean tribes (e.g., Euneomyini, Andinomyini, and Chinchillulini). Additionally, *Delomys* presents a unique combination of morphological features including the presence of low-crowned molars with well-developed lophs, wide interorbital regions, and short palate with short incisive foramina, among other characteristics [98, 99]. This genus has never been associated with Phyllotini taxonomically and including it within the tribe would greatly diminish any morphological construct for the tribe. Therefore, we agree with Pardiñas et al. [87] that *Delomys* is better suited for a separate tribal designation, rather than being folded into Phyllotini.

> Tribe Delomyini, new tribe

> Type genus: *Delomys* Thomas, 1917

> Contents: *Delomys altimontanus* Gonçalves and Oliveira, 2014; *Delomys dorsalis* (Hensel, 1872); *Delomys sublineatus* (Thomas, 1903).

Definition: this new tribe of subfamily Sigmodontinae is nested within a clade containing the tribes Andinomyini, Euneomyini, Wiedomyini, Abrotrichini, and Phyllotini. Delomyini is sister to the tribe Phyllotini, a relationship with high support on the genomic analysis here presented.

Diagnosis: this tribe is diagnosed by the unique combination of: a unique pelage color pattern for the subfamily, with a dark median stripe on dorsal region (darker in *D. sublineatus*, more subtle in *D. altimontanus* and *D. dorsalis*); pelage harsh to dense; mystacial vibrissae long and moderately dense; manus and pes densely covered by white hairs, digits with short ungueal tufts; tail similar to head and body length, moderately to strongly bicolored dorsoventrally; skull robust, with long and narrow rostrum, with distinct rostral tube, frontal sinuses not inflated; interorbital region short and wide, hourglass shaped, with rounded supraorbital margins; fronto-parietal suture wide, concave; interparietal small; alisphenoid strut absent, buccinator-masticatory foramen and accessory oval foramen confluent; anterior opening of the alisphenoid canal open; stapedial foramen, squamoso-alisphenoid groove and sphenofrontal foramen present; posterior opening of alisphenoid canal large and conspicuous, with deep posterior groove or depression; secondary anastomosis of internal carotid artery not visible on dorsal surface of parapterygoid plate; incisive foramina very wide and long, not extending to the anterocones of M1; mesopterygoid fossa equal to parapterygoid fossae; mesopterygoid fossa extending anteriorly to maxillaries, reaching alveolus of molars; anterior margin squared or rounded; upper incisors narrow, with distinctive orange enamel band; opisthodont; maxillary toothrows parallel; labial and lingual flexi of upper molars penetrating at molar midline; molar series bunodont and lophodont; lingual cusps parallel to the labial cusps; molars pentalophodont, complex, with deep labial and lingual flexi, with labial and lingual cingula or styles; first upper molar with deep anteromedian flexus, 13 thoracic and 6 lumbar vertebrae (see [98, 99] for additional information).

Known collection localities: the genus is widely distributed along the Atlantic Forest, occurring from the Brazilian state of Espírito Santo to the Argentinean province of Misiones, from the sea level to mountain regions around 2500 m of elevation [99, 100].

#### Revised Classification

Here we integrate our results and recommendations in a revision of the classification for the subfamily (Table 1).

**Table 1.** A revised and comprehensive classification of the extant Sigmodontinae.

| Higher Tax | Tribes | Subtribes/clades | Genera |
| --- | --- | --- | --- |
| Sigmodontalia | Sigmodontini |  | *Sigmodon* |
|  | Ichthyomyini | Anotomyina | *Anotomys* |
|  |  | Ichthyomyina | *Chibchanomys* |
|  |  |  | *Daptomys* |
|  |  |  | *Ichthyomys* |
|  |  |  | *Neusticomys* |
|  |  |  | *Rheomys* |
| Oryzomyalia |  |  |  |
|  | Oryzomyini | Clade A1 | *Zygodontomys* |
|  |  | Clade A2 | *Scolomys* |
|  |  | Clade B | *Euryoryzomys* |
|  |  |  | *Handleyomys* |
|  |  |  | *Hylaeamys* |
|  |  |  | *Mindomys* |
|  |  |  | *Nephelomys* |
|  |  |  | *Oecomys* |
|  |  |  | *Pattonimus* |
|  |  |  | *Transandinomys* |
|  |  | Clade C | *Microakodontomys* |
|  |  |  | *Microryzomys* |
|  |  |  | *Neacomys* |
|  |  |  | *Oligoryzomys* |
|  |  |  | *Oreoryzomys* |
|  |  | Clade D | *Aegialomys* |
|  |  |  | *Amphinectomys* |
|  |  |  | *Cerradomys* |
|  |  |  | *Drymoreomys* |
|  |  |  | *Eremoryzomys* |
|  |  |  | *Holochilus* |
|  |  |  | *Lundomys* |
|  |  |  | *Melanomys* |
|  |  |  | *Nectomys* |
|  |  |  | *Nesoryzomys* |
|  |  |  | *Oryzomys* |
|  |  |  | *Pseudoryzomys* |
|  |  |  | *Sigmodontomys* |
|  |  |  | *Sooretamys* |
|  |  |  | *Tanyuromys* |
|  | Thomasomyini | Rhagomyina, new rank | *Rhagomys* |
|  |  | Thomasomyina | *Aepeomys* |
|  |  |  | *Chilomys* |
|  |  |  | *Rhipidomys* |
|  |  |  | *Thomasomys* |
|  | Reithrodontini |  | *Reithrodon* |
|  | Akodontini |  | *Abrawayaomys* |
|  |  |  | *Akodon* |
|  |  |  | *Bibimys* |
|  |  |  | *Blarinomys* |
|  |  |  | *Brucepattersonius* |
|  |  |  | *Castoria* |
|  |  |  | *Deltamys* |
|  |  |  | *Gyldenstolpia* |
|  |  |  | *Juscelinomys* |
|  |  |  | *Kunsia* |
|  |  |  | *Lenoxus* |
|  |  |  | *Necromys* |
|  |  |  | *Oxymycterus* |
|  |  |  | *Podoxymys* |
|  |  |  | *Scapteromys* |
|  |  |  | *Thalpomys* |
|  |  |  | *Thaptomys* |
|  | Andinomyini |  | *Andinomys* |
|  |  |  | *Punomys* |
|  | Chinchillulini nov. tax. |  | *Chinchillula* |
|  | Euneomyini |  | *Euneomys* |
|  |  |  | *Irenomys* |
|  |  |  | *Neotomys* |
|  | Neomicroxini |  | *Neomicroxus* |
|  | Wiedomyini |  | *Juliomys* |
|  |  |  | *Phaenomys* |
|  |  |  | *Wiedomys* |
|  |  |  | *Wilfredomys* |
|  | Abrothrichini |  | *Abrothrix* |
|  |  |  | *Chelemys* |
|  |  |  | *Geoxus* |
|  |  |  | *Notiomys* |
|  |  |  | *Paynomys* |
|  |  |  | *Pearsonomys* |
|  | Delomyini nov. tax. |  | *Delomys* |
|  | Phyllotini |  | *Andalgalomys* |
|  |  |  | *Auliscomys* |
|  |  |  | *Calassomys* |
|  |  |  | *Calomys* |
|  |  |  | *Eligmodontia* |
|  |  |  | *Galenomys* |
|  |  |  | *Graomys* |
|  |  |  | *Loxodontomys* |
|  |  |  | *Phyllotis* |
|  |  |  | *Salinomys* |
|  |  |  | *Tapecomys* |

Higher taxa are ordered by the tree in Figure 1 and genera are ordered alphabetically within their containing clades.

## Acknowledgements

Sincere thanks to United States National Museum, University of Michigan Museum of Zoology, University of Washington Burke Museum, Louisiana State University Museum of Zoology, Museu de Zoologia da Universidade de São Paulo, Instituto de Biociências da Universidade de São Paulo and their staffs, and to Carl Saltzberg for providing samples for this study. Analytical resources were provided by the Arkansas High Performance Computing Center (AHPCC).

## Supporting Information

**S1 Fig. Fossil calibrated phylogeny generated using full maximum likelihood in MEGA with dates labeled.** Red diamonds represent calibration points.

**S1 Table. List of samples used, including subfamily and tribal classification, museum voucher number, and biogeographic regions.** Samples from Bangs and Steppan 2022 and Percequillo et al. 2021 are indicated as well. Sequences retrieved from published genomes are listed as “NCBI” under the museum voucher number.

## Notes

### Competing Interest Statement

The authors have declared no competing interest.

## References

1. Schluter D. The ecology of adaptive radiation. Oxford: Oxford University Press; 2000.

2. Losos JB, Ricklefs RE. Adaptation and diversification on islands. Nature. 2009;457:830–6.

3. Patton JL, Pardiñas UFJ, D’Elia G, editors. Mammals of South America. Chicago: University of Chicago Press; 2015.

4. Musser GM, Carleton MD. Superfamily Muroidea. In: Wilson DE, Reeder DM, editors. Mammal species of the world: a taxonomic and geographic reference. 3rd ed. Washington: Smithsonian Institution; 2005.

5. Mammal_Diversity_Database. Mammal Diversity Database. 1.12 ed: Zenodo; 2024.

6. Burgin CJ, Colella JP, Kahn PL, Upham NS. How many species of mammals are there? Journal of Mammalogy. 2018;99:1–14.

7. Steppan SJ, Adkins RM, Anderson J. Phylogeny and divergence-date estimates of rapid radiations in muroid rodents based on multiple nuclear genes. Systematic Biology. 2004;53(4):533–53.

8. Schenk JJ, Rowe KC, Steppan SJ. Ecological opportunity and incumbency in the diversification of repeated continental colonizations by muroid rodents. Systematic Biology. 2013;62(6):837–64.

9. Leite RN, Kolokotronis SO, Almeida FC, Werneck FP, Rogers DS, Weksler M. In the wake of invasion: tracing the historical biogeography of the South American cricetid radiation (Rodentia, Sigmodontinae). PloS one. 2014:9:e100687.

10. Maestri R, Monteiro LR, Fornell R, Upham NS, Patterson BD, de Freitas TRO. The ecology of a continental evolutionary radiation: Is the radiation of sigmodontine rodents adaptive? Evolution. 2017;71(3):610–32.

11. Schenk JJ, Steppan SJ. The role of biogeography in adaptive radiation. American Naturalist. 2018;192:415–31.

12. D’Elía G, Pardiñas UFJ, Patton JL. Subfamily Sigmodontinae Wagner, 1843. Mammals of South America. Chicago: University of Chicago Press; 2015. p. 63–70.

13. Parada A, Hanson J, D’Eiía G. Ultraconserved Elements Improve the Resolution of Difficult Nodes within the Rapid Radiation of Neotropical Sigmodontine Rodents (Cricetidae: Sigmodontinae). Systematic Biology. 2021;70(6):1090–100.

14. Simpson GG. Splendid isolation. The curious history of South American mammals. New Haven: Yale University Press; 1980. 266 p.

15. Reig OA. Diversity patterns and differentiation of high Andean rodents. In: Vuilleumier F, Monasterio M, editors. High altitude tropical biogeography. London: Oxford University Press; 1986. p. 404–39.

16. Smith MF, Patton JL. Phylogenetic relationships and the radiation of sigmodontine rodents in South America: evidence from cytochrome *b*. Journal of Mammalian Evolution. 1999;6(2):89–128.

17. Steppan SJ, Schenk JJ. Muroid rodent phylogenetics: 900-species tree reveals increasing diversification rates. PloS one. 2017:12:e0183070.

18. Vallejos-Garrido P, Pino K, Espinoza-Aravena N, Pari A, Inostroza-Michael O, Toledo-Munoz M, et al. The importance of the Andes in the evolutionary radiation of Sigmodontinae (Rodentia, Cricetidae), the most diverse group of mammals in the Neotropics. Scientific Reports. 2023;13(1):2207.

19. Carleton MD. Phylogenetic relationships on neotomine-peromyscine rodents (Muroidea) and a reappraisal of the dichotomy within New World Cricetinae. Miscellaneous Publications Museum of Zoology, University of Michigan. 1980;157:1–146.

20. Reig OA. A new fossil genus of South American cricetid rodents allied to *Wiedomys*, with an assessment of the Sigmodontinae. Journal of Zoology (London). 1980;192:257–81.

21. Reig OA. Distribuição geográfica e história evolutiva dos roedores muroides sulamericanos (Cricetidae: Sigmodontinae). Rev Brasil Genet. 1984;7:333–65.

22. Steppan SJ. Revision of the tribe Phyllotini (Rodentia: Sigmodontinae), with a phylogenetic hypothesis for the Sigmodontinae. Fieldiana: Zoology, ns. 1995;80:1–112.

23. Vorontsov NN. The system of hamster (Cricetinae) in the sphere of the world fauna and their phylogenetic relations [in Russian]. Biuletin Moskovkogo Obshtschestva Ispitately Prirody, Otdel Biologia. 1959;64:134–7.

24. D’Elîa G. Phylogenetics of sigmodontinae (Rodentia, Muroidea, Cricetidae), with special reference to the akodont group, and with additional comments on historical biogeography. Cladistics. 2003;19(4):307–23.

25. D’Elîa G, Luna L, Gonzalez EM, Patterson BD. On the Sigmodontinae radiation (Rodentia, Cricetidae): An appraisal of the phylogenetic position of Rhagomys. Molecular Phylogenetics and Evolution. 2006;38(2):558–64.

26. Engel SR, Hogan KM, Taylor JF, Davis SK. Molecular systematics and paleobiogeography of the South American sigmodontine rodents. Molecular Biology and Evolution. 1998;15(1):35–49.

27. Parada A, D’Elia G, Palma RE. The influence of ecological and geographical context in the radiation of Neotropical sigmodontine rodents. BMC Evol Biol. 2015;15:172.

28. Parada A, Pardinas UFJ, Salazar-Bravo J, D’Elia G, Palma RE. Dating an impressive Neotropical radiation:. Molecular time estimates for the Sigmodontinae (Rodentia) provide insights into its historical biogeography. Molecular Phylogenetics and Evolution. 2013;66(3):960–8.

29. Weksler M. Phylogeny of Neotropical oryzomyine rodents (Muridae: Sigmodontinae) based on the nuclear IRBP exon. Molecular Phylogenetics and Evolution. 2003;29(2):331–49.

30. Carrizo LV, Tulli MJ, Abdala V. An ecomorphological analysis of forelimb musculotendinous system in sigmodontine rodents (Rodentia, Cricetidae, Sigmodontinae). Journal of Mammalogy. 2014;95(4):843–54.

31. Maestri R, Patterson BD, Fornell R, Monteiro LR, de Freitas TRO. Diet, bite force and skull morphology in the generalist rodent morphotype. Journal of Evolutionary Biology. 2016;29:2191–204.

32. Martínez JJ, Ferro LI, Mollerach MI, Barquez RM. The phylogenetic relationships of the Andean swamp rat genus *Neotomys* (Rodentia, Cricetidae, Sigmodontinae) based on mitochondrial and nuclear markers. Acta Theriol. 2012;57(3):277–87.

33. Salazar-Bravo J, Pardinas UFJ, D’Elia G. A phylogenetic appraisal of Sigmodontinae (Rodentia, Cricetidae) with emphasis on phyllotine genera: systematics and biogeography. Zoologica Scripta. 2013;42(3):250–61.

34. Vilela JF, Mello B, Voloch CM, Schrago CG. Sigmodontine rodents diversified in South America prior to the complete rise of the Panamanian Isthmus. Journal of Zoological Systematics and Evolutionary Research. 2013;52(3):249–56.

35. Gonçalves PR, Christoff AU, Machado LF, Bonvicino CR, Peters FB, Percequillo AR. Unraveling deep branches of the Sigmodontinae tree (Rodentia: Cricetidae) in eastern South America. Journal of Mammalian Evolution. 2020;27(1):139–60.

36. Carrapa B, Clementz M, Feng R. Ecological and hydroclimate responses to strengthening of the Hadley circulation in South America during the Late Miocene cooling. Proc Natl Acad Sci U S A. 2019;116(20):9747–52.

37. Hoorn C, Palazzesi L, Silvestro D. Editorial Preface to Special Issue: Exploring the impact of Andean uplift and climate on life evolution and landscape modification: From Amazonia to Patagonia. Glob Planet Change. 2022;211:6.

38. Hoorn C, Wesselingh FP, ter Steege H, Bermudez MA, Mora A, Sevink J, et al. Amazonia through time: Andean uplift, climate change, landscape evolution, and biodiversity. Science. 2010;330(6006):927–31.

39. Pardiñas UFJ, Curay J, Brito J, Cañón C. A unique cricetid experiment in the northern high-Andean Paramos deserves tribal recognition. Journal of Mammalogy. 2021;102(1):155–72.

40. Bangs MR, Steppan SJ. A rodent anchored hybrid enrichment probe set for a range of phylogenetic utility: From order to species. Mol Ecol Resour. 2022;22(4):1521–8.

41. Percequillo AR, do Prado JR, Abreu EF, Dalapicolla J, Pavan AC, Chiquito ED, et al. Tempo and mode of evolution of oryzomyine rodents (Rodentia, Cricetidae, Sigmodontinae): A phylogenomic approach. Molecular Phylogenetics and Evolution. 2021;159:15.

42. Andermann T, Cano A, Zizka A, Bacon C, Antonelli A. SECAPR-a bioinformatics pipeline for the rapid and user-friendly processing of targeted enriched Illumina sequences, from raw reads to alignments. Peerj. 2018;6:15.

43. Bolger AM, Lohse M, Usadel B. Trimmomatic: a flexible trimmer for Illumina sequence data. Bioinformatics. 2014;30(15):2114–20.

44. Simpson JT, Wong K, Jackman SD, Schein JE, Jones SJM, Birol I. ABySS: A parallel assembler for short read sequence data. Genome Res. 2009;19(6):1117–23.

45. Edgar RC. MUSCLE: multiple sequence alignment with high accuracy and high throughput. Nucleic Acids Res. 2004;32(5):1792–7.

46. Nguyen LT, Schmidt HA, von Haeseler A, Minh BQ. IQ-TREE: A fast and effective stochastic algorithm for estimating maximum-likelihood phylogenies. Molecular Biology and Evolution. 2015;32(1):268–74.

47. Zhang C, Rabiee M, Sayyari E, Mirarab S. ASTRAL-III: polynomial time species tree reconstruction from partially resolved gene trees. BMC Bioinformatics. 2018;19:16.

48. Hoang DT, Chernomor O, von Haeseler A, Minh BQ, Vinh LS. UFBoot2: Improving the Ultrafast Bootstrap Approximation. Molecular Biology and Evolution. 2018;35(2):518–22.

49. Miller MA, Pfeiffer W, T. S. Creating the CIPRES Science Gateway for inference of large phylo-genetic trees. GCE. 2010:1–8.

50. Stamatakis A. RAxML Version 8: A tool for phylogenetic analysis and post-analysis of large phylogenies. Bioinformatics. 2014;30(9):1312–3.

51. Kimura Y, Hawkins MTR, McDonough MM, Jacobs LL, Flynn LJ. Corrected placement of *Mus-Rattus* fossil calibration forces precision in the molecular tree of rodents. Scientific Reports. 2015;5:9.

52. Barbière F, Ortiz PE, Pardiñas UFJ. The oldest sigmodontine rodent revisited and the age of the first South American cricetids. Journal of Paleontology. 2019;93(2):368–84.

53. Smith SA, Brown JW, Walker JF. So many genes, so little time: A practical approach to divergence-time estimation in the genomic era. PloS one. 2018;13(5):18.

54. IUCN. The IUCN Red List of Threatened Species. Version 2021-3 2021 [Available from: https://www.iucnredlist.org.

55. Griffith GE, Omernik JM, Azevedo SH. Ecological classification of the Western Hemisphere. . In: U.S. Environmental Protection Agency WED, editor. Corvallis, OR. 1998. p. 49p. .

56. Yu Y, Harris AJ, Blair C, He XJ. RASP (Reconstruct Ancestral State in Phylogenies): A tool for historical biogeography. Molecular Phylogenetics and Evolution. 2015;87:46–9.

57. Felsenstein J. Evolutionary trees from dna-sequences: a maximum-likelihood approach. Journal of Molecular Evolution. 1981;17(6):368–76.

58. Wilson DE, Reeder DM, editors. Mammal species of the world: a taxonomic and geographic reference (Vol. 1). : JHU Press; 2005.

59. Hershkovitz P. Mice, land bridges and Latin American faunal interchange. In: Wenzel RL, Tipton VJ, editors. Ectoparasites of Panama. Chicago: Field Museum of Natural History; 1966. p. 725–51.

60. Hershkovitz P. The Recent mammals of the Neotropical region: a zoogeographic and ecologic review. In: Keast A, Erk FC, Glass B, editors. Evolution, mammals, and southern continents. Albany: State University of New York; 1972. p. 311–431.

61. Jacobs LL, Lindsay EH. Holarctic radiation of Neogene muroid rodents and the origin of South American cricetids. Journal of Vertebrate Paleontology. 1984;4(2):265–72.

62. Marshall LG. A model for paleobiogeography of South American cricetine rodents. Paleobiology. 1979;5:126–32.

63. Salazar-Bravo J, Tinoco N, Zeballos H, Brito J, Arenas-Viveros D, Marin CD, et al. Systematics and diversification of the Ichthyomyini (Cricetidae, Sigmodontinae) revisited: evidence from molecular, morphological, and combined approaches. PeerJ. 2023;11:e14319.

64. Carranza-Castañeda O, Wolton AH. Cricetid rodents from the Rancho El Ocote fauna, late Hemphillian (Pliocene), Guanajuato. Rev Mex de Cien Geol. 1992;10:71–93.

65. Czaplewski NJ. Sigmodont rodents (Mammalia; Muroidea; Sigmodontinae) from the Pliocene (early Blancan) Verde Formation, Arizona. Journal of Vertebrate Paleontology. 1987;7:183–99.

66. Jacobs LL, Lindsay EH. *Prosigmodon oroscoi*, a new sigmodont rodent from the late Tertiary of Mexico. Journal of Paleontology. 1981;55:425–30.

67. Reig OA. Roedores cricétidos del Plioceno superior de la Provincia de Buenos Aires. Publnes Mus munic Cien nat "Lorenzo Scaglia". 1978;2(8):164–90.

68. Pardiñas UFJ, D’Elia GD, Ortiz PE. Sigmodontinos fósiles (Rodentia, Muroidea, Sigmodontinae) de América del Sur: estadoactual de su conocimiento y prospectiva. Mast Neot. 2002;9(2):209–52.

69. Garralla SS, Anzótegui LM, Mautino LR. Relaciones paleoflorísticas del Mioceno-Plioceno del norte argentino2016; 16(1).

70. Prevosti FJ, Romano CO, Forasiepi AM, Hemming S, Bonini R, Candela AM, et al. New radiometric (40)Ar-(39)Ar dates and faunistic analyses refine evolutionary dynamics of Neogene vertebrate assemblages in southern South America. Scientific Reports. 2021;11(1):9830.

71. Sanín MJ, Mejía-Franco FG, Paris M, Valencia-Montoya WA, Salamin N, Kessler M, et al. Geogenomics of montane palms points to Miocene-Pliocene Andean segmentation related to strike-slip tectonics. Journal of Biogeography. 2022;49(9):1711–25.

72. Chaves JA, Weir JT, Smith TB. Diversification in *Adelomyia* hummingbirds follows Andean uplift. Molecular Ecology. 2011;20(21):4564–76.

73. Vieu JC, Hughes CE, Kissling J, Grant JR. Evolutionary diversification in the hyper-diverse montane forests of the tropical Andes: radiation of *Macrocarpaea* (Gentianaceae) and the possible role of range expansion. Bot J Linnean Soc. 2022;199(1):53–75.

74. Graham A. Studies in neotropical paleobotany. XIII. An Oligo-Miocene palynoflora from Simojovel (Chiapas, Mexico). Am J Bot. 1999;86(1):17–31.

75. Ornelas JF, Ruiz-Sánchez E, Sosa V. Phylogeography of *Podocarpus matudae* (Podocarpaceae): pre-Quaternary relicts in northern Mesoamerican cloud forests. Journal of Biogeography. 2010;37(12):2384–96.

76. Palacios-Chávez R, Rzedowski J. Estudio palinologico de las floras fosiles del Mioceno inferior y principios del Mioceno de la región de Pichucalco, Chiapas, Mexico. Acta Bot Mex. 1993;24:1–96.

77. Piperno DR, Moreno JE, Iriarte J, Hoist I, Lachniet M, Jones JG, et al. Late pleistocene and holocene environmental history of the iguala valley, central balsas watershed of Mexico. Proc Natl Acad Sci U S A. 2007;104(29):11874–81.

78. Vizcaíno SF, Fariña RA, Zárate MA, Bargo MS, Schultz P. Palaeoecological implications of the mid-Pliocene faunal turnover in the Pampean Region (Argentina). Paleogeogr Paleoclimatol Paleoecol. 2004;213(1-2):101–13.

79. Wagner JA. Die säugethiere in Abbildungen nach der nature mit Beschreibung von Dr. Johann Christian Daniel von Schreber. Suppl. 3: Erlangen (Germany); 1843. 614 p.

80. Cockerell TDA, Miller LI, Printz M. The auditory ossicles of American rodents. Bulletin of the American Museum of Natural History. 1914;33:347–64.

81. Vorontsov NN. Sistema khomiakov (Cricetinae) mirovoi fauny i ikh filogeneticheskie sviazi. Biuletin Moskovkogo Obshtschestva Ispitately Prirody, Otdel Biologia. 1959; 44:134–7.

82. Steadman DW, Ray C, E. Relationships of *Megaoryzomys curioi*, an extinct cricetine rodent (Muroidea: Muridae) from the Galápagos Islands, Ecuador Smithsonian Contributions to Paleobiology. 1982;51.

83. Pardiñas UFJ, Teta P, Salazar-Bravo J. Una nueva tribu de roedores Sigmodontinae (Cricetidae). Mast Neot. 2015;22:171–87.

84. Salazar Bravo J, Pardiñas UFJ, Zeballos H, Teta P. Description of a new tribe of sigmodontine rodents (Cricetidae: Sigmodontinae) with an updated summary of valid tribes and their generic contents. Occ Paper Mus Texas Tech. 2016;338.

85. Cazzaniga NJ, Cañon CA, Pardiñas UFJ. The availability, authorships and dates of tribal names in the Sigmodontinae (Rodentia, Cricetidae) current classification. Bionomina. 2019;15:37–50.

86. ICZN. International code of zoological nomenclature. Fourth ed. London: The International Trust for Zoological Nomenclature; 1999. 306 p.

87. Pardinas UFJ, Tinoco N, Barbière F, Ronez C, Cañón C, Lessa G, et al. Morphological disparity in a hyperdiverse mammal clade: a new morphotype and tribe of Neotropical cricetids. Zool J Linn Soc. 2022;196(3):1013–38.

88. Braun JK. Systematic relationships of the tribe Phyllotini (Muridae: Sigmodontinae) of South America. Norman: Special Publication, Oklahoma Museum of Natural History; 1993. 50 p.

89. Hershkovitz P. Evolution of Neotropical cricetine rodents (Muridae) with special reference to the phyllotine group. Fieldiana: Zool. 1962;46:1–524.

90. Olds N, Anderson S. A diagnosis of the tribe Phyllotini (Rodentia, Muridae). In: Redford KH, Eisenberg JF, editors. Advances in Neotropical mammalogy. Gainesville, Florida: Sandhill Crane Press; 1989. p. 55–74.

91. Jansa SA, Weksler M. Phylogeny of muroid rodents: relationships within and among major lineages as determined by IRBP gene sequences. Molecular Phylogenetics and Evolution. 2004;31(1):256–76.

92. Anderson S, Yates TL. A new genus and species of phyllotine rodent from Bolivia. Journal of Mammalogy. 2000;81(1):18–36.

93. Pacheco V, Cairampoma R, Quispe A, Velezvilla G, Vasquez A. Notable range extension *Chinchillula sahamae* Thomas, 1898 (Rodentia, Sigmodontinae) to central Peru, with natural history notes. Check List. 2023;19(2):183–90.

94. Salazar-Bravo J. Genus *Chinchillula* Thomas, 1898. In: Patton JL, Pardiñas UFJ, D’Elía G, editors. Mammals of South America: Rodents. 2. Chicago, USA: University of Chicago Press; 2015. p. 78–9.

95. Stephens L, Traylor MA. Ornithological Gazetteer of Peru: Bird Department, Museum of Comparative Zoology, Harvard University; 1983.

96. Percequillo AR, Braga CAD, Brandao MV, de Abreu EF, Gualda-Barros J, Lessa GM, et al. The genus *Abrawayaomys* Cunha and Cruz, 1979 (Rodentia: Cricetidae: Sigmodontinae): geographic variation and species definition. Journal of Mammalogy. 2017;98(2):438–55.

97. Pardiñas UFJ, Teta P, D’Elía G. Taxonomy and distribution of *Abrawayaomys* (Rodentia: Cricetidae), an Atlantic Forest endemic with the description of a new species. Zootaxa. 2009(2128):39–60.

98. Voss RS. A revision of the Brazilian muroid rodent genus *Delomys* with remarks on "thomasomyine" characters. American Museum Novitates. 1993;3073:1–46.

99. Voss RS. Genus *Delomys* Thomas, 1917. In: Patton JL, Pardiñas UFJ, D’Elía G, editors. Mammals of South America: Rodents. 2. Chicago, USA: University of Chicago Press; 2015. p. 79–83.

100. Gonçalves PR, de Olivera JA. An integrative appraisal of the diversification in the Atlantic forest genus *Delomys* (Rodentia: Cricetidae: Sigmodontinae) with the description of a new species. Zootaxa. 2014;3760(1):001–38.

